# Electrical perturbation reveals a directional architecture for communication across human visual streams

**DOI:** 10.64898/2026.04.14.718565

**Authors:** María Guadalupe Yáñez-Ramos, Gabriela Ojeda Valencia, Harvey Huang, Nicholas M. Gregg, Jordan Bilderbeek, Morgan Montoya, Kendrick Kay, Gregory A. Worrell, Kai J. Miller, Dora Hermes

## Abstract

Human visual cortex comprises three interacting streams, but anatomy and correlated activity cannot determine the direction or reliability of interareal influence. Characterizing directional interactions among these streams can reveal the architecture through which activity can propagate across the visual system. Here, we used single-pulse electrical stimulation during intracranial EEG recordings in 23 patients to map directed effective connectivity among 22 atlas-defined visual cortical areas. The resulting effective connectivity matrix revealed a selective and asymmetric architecture of interareal influence. Feedforward influences from early visual areas to the dorsal and lateral streams were more reliable and more prevalent than the corresponding feedback influences, whereas interactions between early visual and ventral temporal areas were comparatively balanced. Cross-stream interactions favored temporal-to-parietal over parietal-to-temporal influence, with higher response reliability and a greater proportion of significant responses among sampled connections. Network-level profiles were consistent with source-like organization in early visual areas and the ventral stream, and integrator-like organization in the dorsal and lateral streams. Visual-task data collected in three of the same patients illustrated how stimulation-derived connectivity may relate to functional responses across connected visual regions. These findings reveal a directional architecture for activity propagation across the human visual cortex and provide empirical constraints for biologically grounded models of visual processing.

## Introduction

The human visual system is organized into three cortical streams that support perception and visually guided action. Classical dual-stream models distinguish a ventral ‘what’ stream, located within the inferior temporal cortex and specialized for object recognition, and a dorsal ‘where/how’ stream, extending through occipitoparietal regions and supporting visuospatial processing and visually guided behavior^1,2^. Recent work identified a third lateral visual stream in the lateral occipitotemporal cortex toward the superior temporal sulcus that is potentially unique to the human brain^2–8^ and is proposed to support dynamic and socially relevant visual representations^9–14^.

Although early theories proposed functionally segregated processing within each stream, more recent models emphasize feedback and continuous cross-talk between streams^15–19^. Consistent with this view, functional interactions between visual streams have been observed during perceptual tasks^20–24^. Such cortico-cortical cross-talk has been proposed to play a central role in visual computation by facilitating prediction and error detection^25,26^. However, correlated activity cannot determine which region drives another, whether influence is reciprocal, or whether an interaction is spatially broad or confined to a subset of source and target sites.

While the flow of information between visual streams is not well understood, some of the anatomical pathways that support these interactions are well described. Human diffusion MRI studies have shown that the vertical occipital fasciculus (VOF) and the IPS-FG (intraparietal sulcus, fusiform gyrus) pathways connect dorsal and ventral streams^27–31^. In non-human primates, retrograde tracing studies have characterized detailed, directed, and weighted connectivity matrices showing multiple connections between dorsal and ventral areas^32,33^. However, whether interactions in human brains are unidirectional or reciprocal, and whether they are spatially specific or broadly distributed, remains unclear. Connectivity profiles can also predict functional specialization within ventral temporal cortex^34^.

The overarching goal of the present study was to characterize the directionality, response reliability, and spatial distribution of interactions between human visual cortical areas. Using single-pulse electrical stimulation (SPES) as a causal perturbation during intracranial stereo EEG recordings (sEEG), we probed interareal influence with millisecond precision in humans^35–37^. Such controlled, artificial perturbations can isolate relationships and identify organizing principles of cortical networks^38^. Previous SPES studies have suggested that this approach can capture a predominance of feedforward signal propagation from early to higher-order visual areas^39^ and clustered influences, for example on the collateral sulcus from limbic and cortical structures^40^. We therefore stimulated electrodes across early visual areas and the three visual streams in 23 patients and measured brain stimulation-evoked potentials (BSEPs) from gray matter sites. We quantified feedforward versus feedback influences between early visual areas and higher-order areas in each visual stream, as well as interactions between streams. We also quantified outgoing and incoming influence profiles, separated spatial influence at the source and target ends, and examined dorsal-stream interactions with A37elv together with visual-task responses in three participants. These additional analyses allowed us to illustrate how SPES-derived connectivity may relate to functional responses measured in visual-task data.

Our results reveal spatially selective cross-stream routing between 22 visual areas, and stream-dependent directional asymmetries that distinguish source-like from integrator-like regions within the sampled human visual cortex. Together, the findings provide a perturbation-based characterization of directional interactions across human visual streams.

## Results

To characterize how visual areas influence one another, we used SPES and sEEG to record BSEPs and measure effective connectivity among visual areas in 23 patients undergoing clinical monitoring for drug-resistant epilepsy. We sampled all possible connections permitted by the available gray matter electrode coverage across early visual areas and the dorsal, lateral, and ventral visual streams. Reliable influences are expected to produce reproducible responses across trials. We therefore quantified directional influence using the coefficient of determination (CoD), an index of cross-trial response reliability. This metric can be directly compared across subjects and sites given its independence from the overall response amplitude. Sampled response prevalence was calculated as the proportion of significant responses among all tested possible connections. This allowed us to assess directional preferences and the prevalence of detectable effective connectivity across the sampled human visual streams. Source and target ratios further quantified the proportion of effective stimulation pairs and responsive recording electrodes.

### Feedforward and feedback influence in individual participants

To test whether early visual areas exert feedforward influences on higher-order visual streams, we delivered SPES through contact pairs located in V1, V2, and V3 and analyzed BSEPs recorded across dorsal, lateral, and ventral streams. Before presenting the group results, we show individual-participant examples illustrating these effects. Fig 1 shows four representative cases (one stimulation pair per participant) in which stimulation of V1-V3 elicited reliable responses in dorsal (IPS), lateral (LO; lateral-occipital cortex), and ventral (FG) areas. Together, these examples indicate that perturbations of a single early visual site can elicit distributed responses across visual cortical areas, consistent with a feedforward organization.

**Fig 1.**
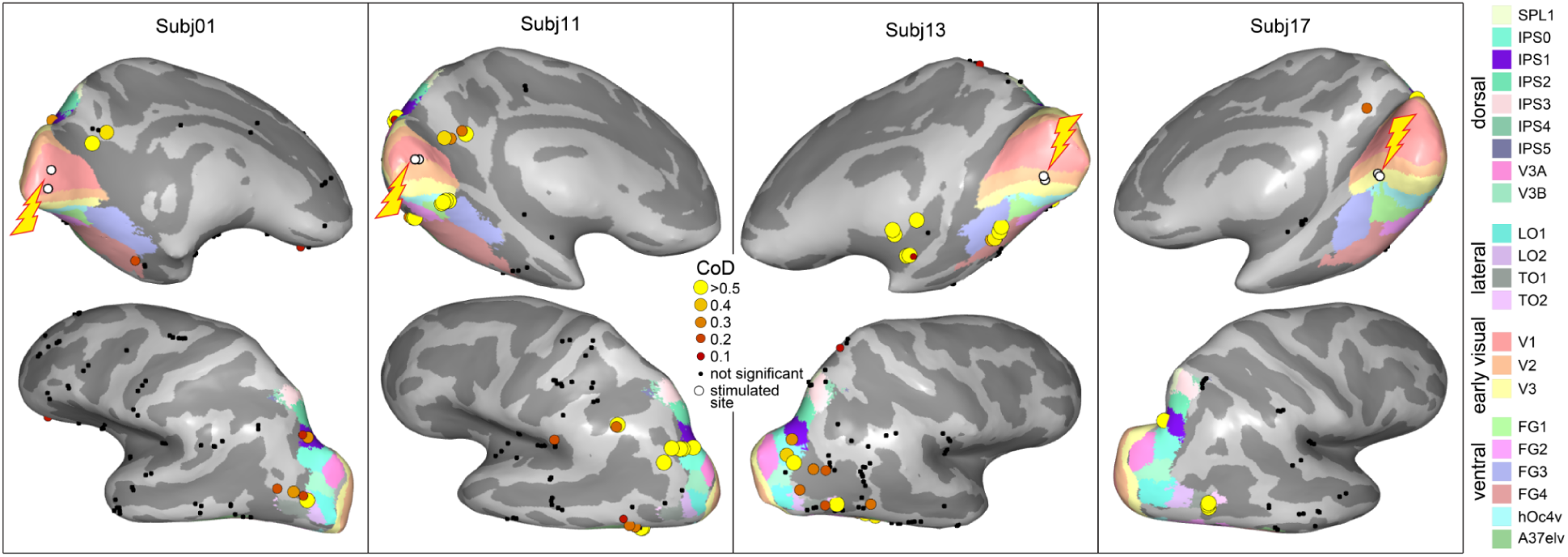
Stimulation of early visual areas evokes feedforward responses across dorsal, lateral, and ventral visual streams. Illustrative examples from four representative participants (subj01, subj11, subj13, subj17) are shown on the inflated cortical surface. Colored dots indicate recording sites with significant brain stimulation-evoked potentials (BSEPs), with color denoting response reliability quantified by the coefficient of determination (CoD). Black dots indicate gray matter electrode contacts without significant responses. The lightning bolt points to the two electrode contacts (white circles) that were stimulated. All gray matter electrode contacts are shown for reference. Subj01: V1 stimulation elicited significant responses in IPS1, LO2, and FG4. Subj11: V1 stimulation elicited significant responses in IPS0, FG1, and FG4. Subj13: V2 stimulation elicited significant responses in IPS0, LO2, TO2, and FG3. Subj17: V3 stimulation elicited significant responses in IPS0 and TO1. Colormaps in the cortical renderings correspond to visual areas defined using probabilistic atlases (see Methods).

To directly compare feedforward and feedback influence, we next examined connections between early visual areas and higher-order regions in two individual examples in which bidirectional testing was feasible using the same four electrodes (Fig 2). In each case, one electrode pair was stimulated while responses were recorded from the second pair, and the direction was then reversed by stimulating the second pair while recording from the first pair. In these examples, stimulation of V2 elicited reliable responses in TO1 (temporal-occipital cortex) of the lateral stream (feedforward direction), whereas stimulation of TO1 did not elicit reliable responses in V2 (feedback direction), illustrating directional asymmetry at the individual level.

**Fig 2.**
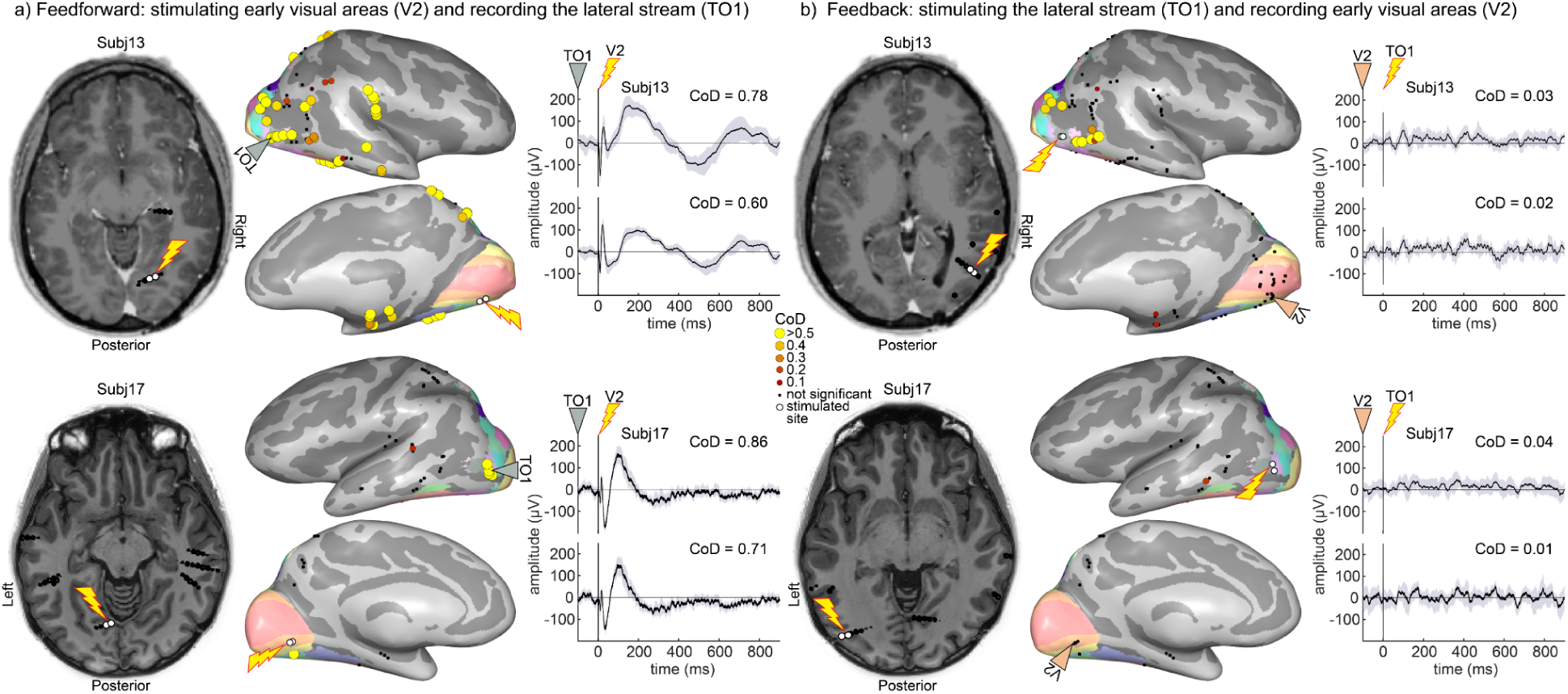
Effective connectivity between the early visual cortex and the lateral stream is asymmetric. Bidirectional testing of effective connectivity between early visual area V2 and the lateral stream (TO1) is shown for two representative participants (subj13, subj17). **a)** Feedforward direction. Stimulation of an electrode pair located in V2 (lightning bolt pointing to white circled electrodes) elicited significant brain stimulation-evoked potentials (BSEPs) in two neighboring electrodes located in TO1 (gray triangle) in subject 13 (top) and subject 17 (bottom). Axial T1-weighted MRI (left) shows the stimulated electrode contact locations. Inflated cortical surfaces (center) show significant response sites colored by the coefficient of determination (CoD). Black dots indicate gray matter contacts without significant responses. Average BSEP traces recorded from the two electrodes in TO1 (right; mean ± 95% CI) are shown, aligned to stimulation onset (0 ms). **b)** Feedback direction. In the same participants, stimulation of the two electrodes in TO1 (lightning bolt pointing to white circled electrodes) did not elicit significant responses in the two electrodes in V2 (orange triangle). Corresponding MRI slices, cortical surface maps, and BSEP traces are shown using the same conventions as in panel a). Colormaps of the renderings are the same as in Fig 1.

Bidirectional testing using the same four electrodes was not always possible because of clinical and methodological constraints, including the requirement that stimulation contacts be located in gray matter. White matter stimulation can result in both orthodromic and antidromic signal propagation^41^, making rigorous gray matter inclusion important for establishing asymmetries (see also S1 Fig.). In the two examples shown here, feedforward responses exhibited high CoD values (0.60-0.86), whereas feedback responses showed low CoD values or were absent (CoD = 0.01-0.04). These individual examples illustrate that effective connectivity between visual areas can be asymmetric, favoring the feedforward direction in these sampled connections.

### Feedforward influences to dorsal and lateral streams are more reliable and prevalent than feedback influences

We next quantified response reliability of feedforward and feedback influences across the group. A total of 1096 possible connections in the feedforward-feedback directions across participants with relevant coverage of early visual areas and at least one higher-order visual streams, were included in this analysis. Linear mixed-effects modeling revealed that feedforward influences were significantly more reliable than feedback influences (Fig 3a, left) and exhibited significantly higher CoD values (β_1_ = 0.132, SE = 0.014, t(1094) = 9.46, p = 1.79e-20, 95% CI [0.105, 0.160], see S1 Table for full results). This feedforward dominance was observed in the dorsal and lateral streams (Fig 3a; dorsal: β_1_ = 0.213, SE = 0.024, t(337) = 8.93, p = 2.73e-17, 95% CI [0.166, 0.259], lateral: β_1_ = 0.191, SE = 0.024, t(309) = 7.92, p = 4.46e-14, 95% CI [0.143, 0.238]). Both comparisons remained significant after false discovery rate correction (dorsal: q = 8.19e-17; lateral: q = 6.69e-14). In contrast response reliability was comparatively balanced between early visual areas and the ventral stream with no significant directional difference (β_1_ = 0.039, SE = 0.021, t(444) = 1.80, p = 0.073, q = 0.073, 95% CI [-0.004, 0.081]). This stream-specific pattern indicates strong feedforward asymmetry for dorsal and lateral streams, but comparatively balanced reliability between early visual and ventral areas.

**Fig 3.**
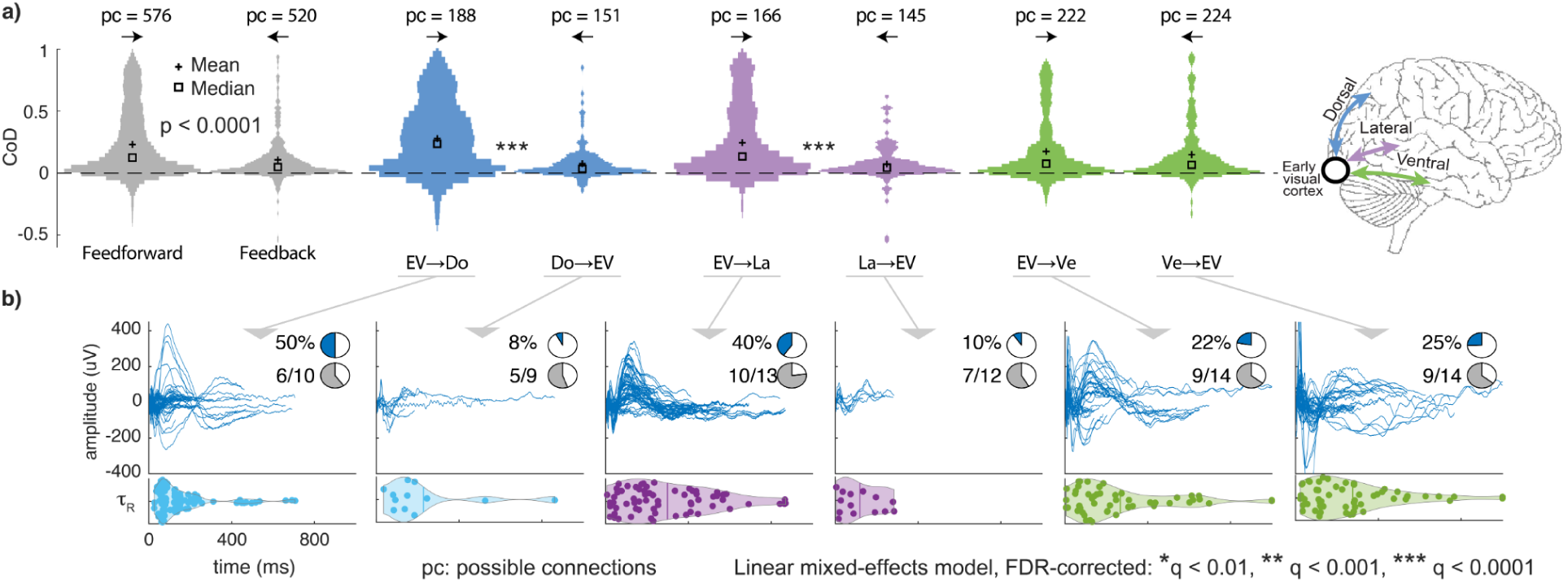
Feedforward influences to the dorsal and lateral streams are more reliable and prevalent than feedback influences. **a)** Violin plots show the distribution of the coefficient of determination (CoD) values, an index of cross-trial response reliability, for feedforward (→) and feedback (←) directions. The left pair (gray) summarizes all connections between early visual areas (EV) and the three visual streams combined. Subsequent pairs show the same comparison separately for the dorsal (Do; blue), lateral (La; lilac), and ventral (Ve; green) streams. Feedforward influences exhibited significantly higher CoD values than feedback influences overall and for the dorsal and lateral streams, but not for the ventral stream. The overall comparison is reported as p; stream-specific pairwise comparisons were false discovery rate corrected (*q < 0.01, **q < 0.001, ***q < 0.0001). pc denotes the number of possible connections tested. **b)** Stimulation-evoked responses across all participants, shown separately for each feedforward (left) and feedback (right) direction. Colored traces in blue represent significant BSEPs. Pie charts summarize the percentage of significant responses (blue) and the proportion of participants showing at least one significant response among those with possible connections (gray). Horizontal violin plots show the distribution of response durations (*τ*_R_); duration comparisons are descriptive.

Analysis of spatial prevalence revealed a corresponding stream dependent pattern of asymmetry. Significant responses were observed in 36.5% of all possible feedforward connections tested (210/576), compared with 16.0% of all possible feedback connections tested (83/520). When separated by stream (Fig 3b), feedforward responses occurred in 50%, 40%, and 22% of possible connections from early visual areas to dorsal, lateral, and ventral streams, respectively, compared to 8%, 10%, and 25% in the corresponding feedback directions. Thus, the dorsal and lateral pathways showed pronounced feedforward asymmetry in response prevalence, whereas interactions between early visual and ventral areas were comparatively balanced. For reference, across all tested interactions between visual areas, 26.24% of possible connections were significant. Thus, feedforward responses were detected across more sampled connections than average, whereas feedback responses were detected across fewer. These percentages describe the extent of detectable influence within sampled contacts.

Response duration distributions (*τ*_R_) are shown descriptively in Fig 3b. Together, these results show a robust but stream-dependent directional asymmetry: feedforward influences were more reliable and prevalent overall, with pronounced asymmetries in dorsal and lateral streams, whereas interactions between early visual and ventral areas were comparatively balanced.

### Temporal-to-parietal influences are overall more reliable and prevalent than parietal-to-temporal influences

We next characterized directional asymmetries between the visual streams. We defined upward influences as ventral → lateral, ventral → dorsal, and lateral → dorsal connections, corresponding to a temporal-to-parietal direction. The three reverse connections were defined as downward influences, corresponding to a parietal-to-temporal direction. These comparisons included 735 possible connections (396 upward and 339 downward) across participants with relevant cross-stream coverage. Linear mixed-effects modeling revealed that, when the three stream pairs were analyzed together, upward influences exhibited significantly higher CoD values than downward influences (Fig 4a, β_1_ = 0.053, SE = 0.016, t(733) = 3.37, p = 7.92e-4, 95% CI [0.022, 0.084]). When stream pairs were examined separately (ventral-lateral, ventral-dorsal, and lateral-dorsal), CoD significantly favored the upward direction for ventral-to-lateral and ventral-to-dorsal influences, but not for lateral-to-dorsal influences (ventral→lateral: β_1_ = 0.076, SE = 0.026, t(281) = 2.90, p = 0.0040, q = 0.0102, 95% CI [0.024, 0.128]; ventral→dorsal: β_1_ = 0.054, SE = 0.020, t(321) = 2.72, p = 0.0068, q = 0.0102, 95% CI [0.015, 0.093]; lateral→dorsal: β_1_ = 0.046, SE = 0.035, t(127) = 1.34, p = 0.184, q = 0.184, 95% CI [-0.022, 0.115]; see S2 Table for full statistics). Thus, the temporal-to-parietal reliability bias was present overall but heterogeneous across stream pairs, with significant directional asymmetries in two of the three comparisons.

**Fig 4.**
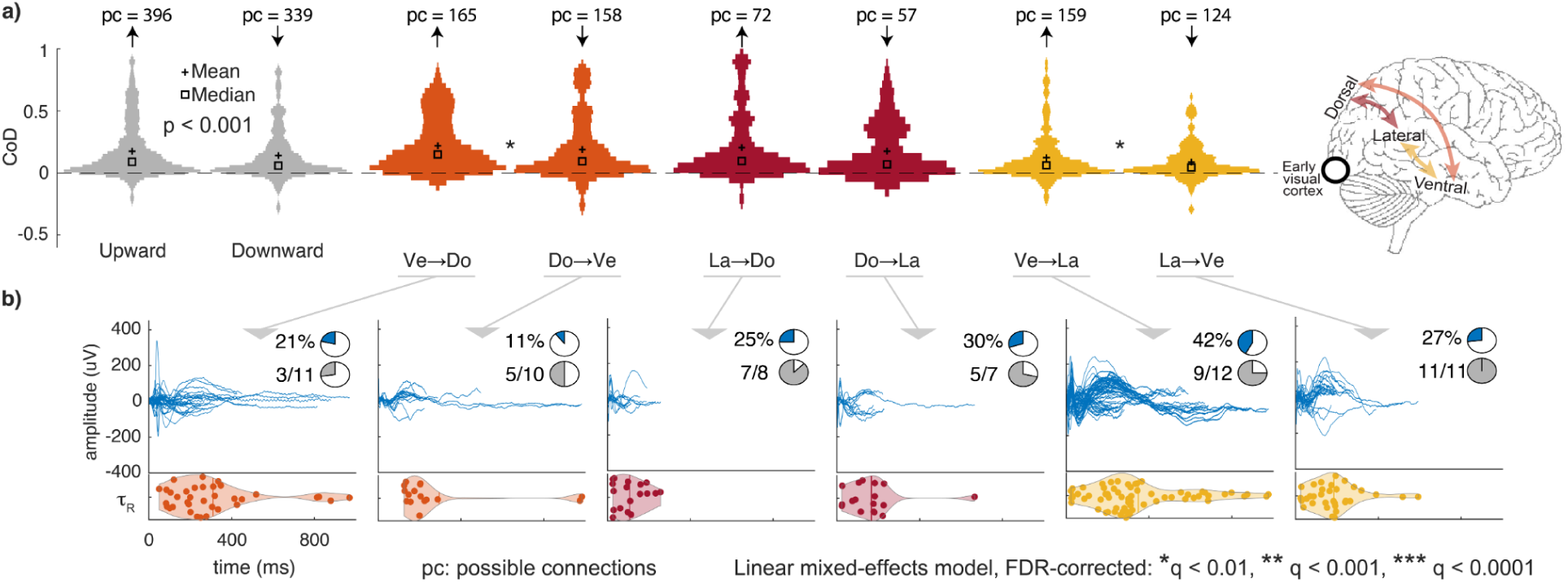
Temporal-to-parietal influences show greater response reliability than parietal-to-temporal influences. **a)** Violin plots show the distribution of the coefficient of determination (CoD) values for upward influences (↑; temporal-to-parietal: Ve→La, Ve→Do, and La→Do) and downward (↓; the corresponding reverse directions) directions, following the same conventions as Fig 3. The left pair (gray) summarizes the three stream pairs combined. Subsequent pairs show the same comparison for individual stream interactions: ventral to dorsal (Ve→Do; orange), lateral to dorsal (La→Do; dark red), and ventral to lateral (Ve→La; yellow). Across all cross-stream connections, temporal-to-parietal influences exhibited significantly higher CoD values than parietal-to-temporal influences (linear mixed-effects model, p < 0.001). Pairwise stream comparisons were FDR-corrected; ventral→dorsal and ventral→lateral were significant, whereas lateral→dorsal was not (*q < 0.01, **q < 0.001, ***q < 0.0001). pc denotes the number of possible connections tested. **b)** Stimulation-evoked responses across all participants, shown separately for each upward (left) and downward (right) direction. Traces represent significant BSEPs. Pie charts summarize the proportion of significant responses (blue) and the fraction of participants showing at least one significant response among those with possible connections (gray). Horizontal violin plots show the distribution of response durations (*τ*_R_); duration comparisons are descriptive.

At the combined level, sampled response prevalence showed the same temporal-to-parietal preference. Significant responses were observed in 30.3% of upward connections (120 of 396), compared with 19.8% of downward connections (67 of 339). When examined separately, significant responses occurred in 21% of ventral→dorsal, 25% of lateral→dorsal, and 42% of ventral→lateral connections, compared with 11%, 30%, and 27% in the corresponding reverse directions, respectively. Thus, the prevalence asymmetry was also stream dependent: it favored the temporal-to-parietal direction for ventral–dorsal and particularly ventral–lateral interactions, whereas lateral–dorsal interactions were comparatively balanced.

Response-duration distributions (*τ*R) are shown descriptively (Fig 4b). Together, these results indicate that cross-stream influence is directionally asymmetric at the group level, with an overall temporal-to-parietal preference that varies across individual stream interactions.

### Output-to-input ratios suggest distinct network profiles

In directed graph theory, the functional role of a node can be inferred from the ratio between incoming and outgoing connections^42^. Nodes with many incoming connections and few outgoing connections function as integrators, whereas nodes with more outgoing than incoming connections function as sources or projectors. At the cortical level, studies using SPES have applied similar logic to infer functional specialization based on patterns of effective connectivity^43,44^. Here, we applied this framework to our SPES measurements to test whether visual regions occupy different organizational profiles within the sampled visual cortical network. We treated early visual areas, dorsal, lateral, and ventral streams as units of a single visual cortical network. Analyses were restricted to interactions within this network. For each unit, we computed the output-to-input ratio, defined as the proportion of significant outgoing responses divided by the proportion of significant incoming responses, providing an index of each unit’s relative influence within the sampled network. Connections involving areas outside the defined visual cortical network were not considered (e.g. connections with frontal eye fields).

Early visual areas exhibited a strong predominance of outgoing influences relative to incoming influences, resulting in a high output-to-input ratio (2.28). This pattern was consistent with higher response reliability for outgoing than incoming influences. This pattern suggests that early visual areas show a source-like profile, exerting influence on dorsal, lateral, and ventral streams within the sampled network. The ventral stream showed a predominance of outgoing over incoming influence (output-to-input ratio = 1.48), suggesting that it also exhibits a source-like profile within the sampled network. In contrast, the lateral and dorsal streams showed fewer outgoing than incoming influences (output-to-input ratios = 0.48 and 0.36, respectively), consistent with integrator-like profiles within the visual cortical network (Fig 5). This pattern was consistent with higher response reliability for temporal-to-parietal than parietal-to-temporal influences. We emphasize that these roles are specific to the visual cortical network examined here and may differ when these regions participate in other cortical networks.

**Fig 5.**
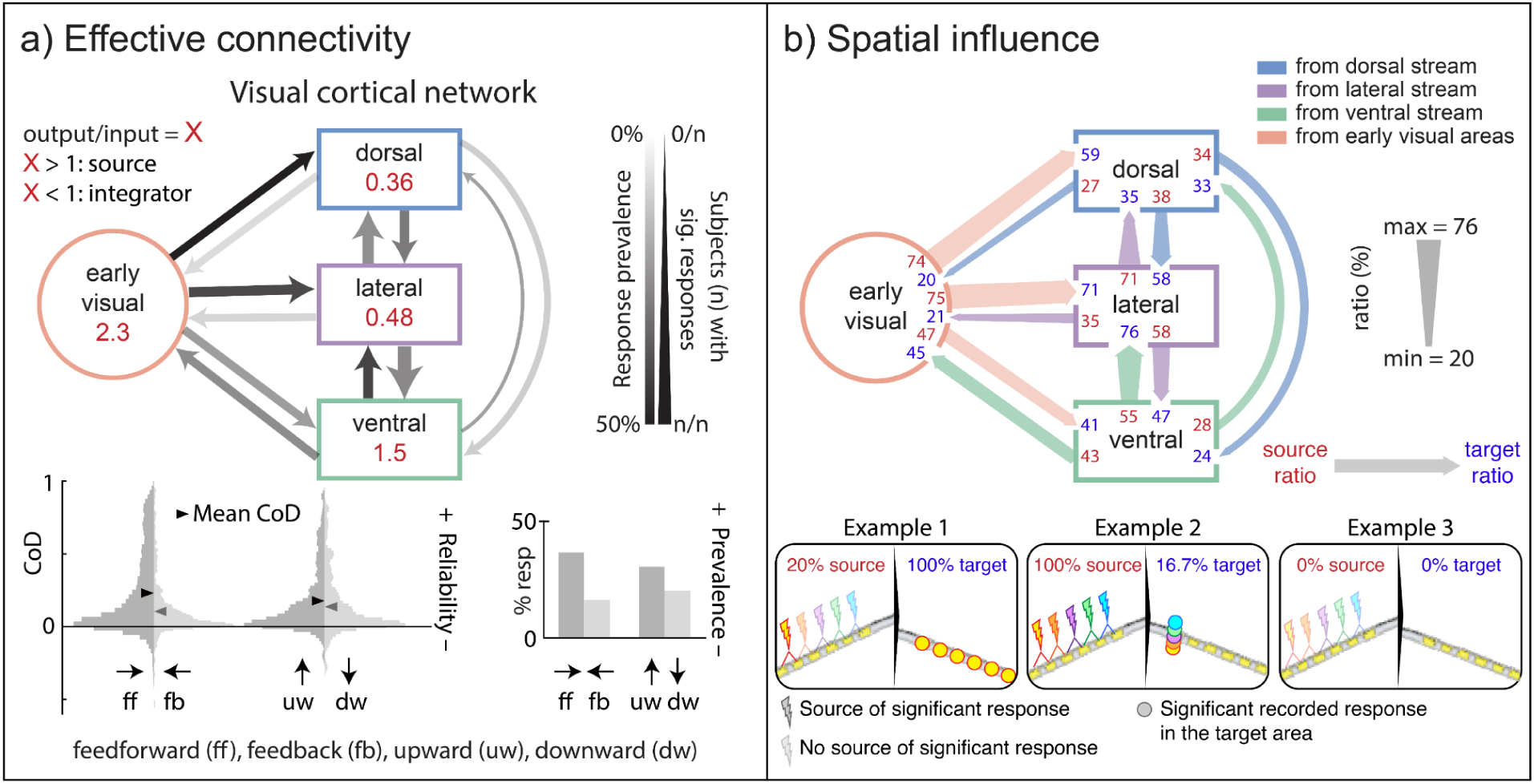
Directional asymmetries and network profiles across the human visual cortical network. This schematic integrates directionality, response reliability, sampled response prevalence, and network profiles derived from effective connectivity measurements across early visual areas, dorsal, lateral, and ventral streams. **a)** Effective connectivity summary. Top: arrows indicate the direction of influence. Arrow opacity is scaled according to the proportion of significant (sig.) responses across all tested connections, whereas arrow thickness represents the proportion of participants showing at least one significant response for that direction among participants with coverage. Red numbers inside each node indicate the output-to-input ratio (X), calculated as the proportion of significant outgoing divided by the proportion of significant incoming responses. Values greater than 1 indicate a source-like profile, whereas values less than 1 indicate an integrator-like profile within the sampled network. Bottom left: distributions of coefficient of determination (CoD) values for feedforward versus feedback influences and upward (temporal-to-parietal) versus downward (parietal-to-temporal) influences; triangles indicate the mean CoD. Bottom right: percentage of significant responses (% resp), with higher percentages indicating greater response prevalence across sampled connections. **b)** Source-side and target-side spatial influence. Arrow color indicates the source region. Tail width and red values indicate the source ratio, defined as the percentage of source stimulation pairs that evoked at least one significant response in the target region. Arrowhead width and blue values indicate the target ratio, defined as the percentage of target recording electrodes that showed at least one significant response following stimulation of the source region. Wider arrow segments indicate broader spatial influence, whereas narrower segments indicate more spatially restricted influence. Example 1 illustrates a source-focal and target-broad interaction; example 2 illustrates a source-broad and target-focal interaction; and example 3 illustrates no detected significant response.

### Area-specific organization of effective connectivity across visual cortex

Each visual stream comprises multiple visual areas that may contribute differently to network organization. Tracer studies in macaques have shown that individual areas within the same stream can exhibit distinct connectivity profiles^32,33^. However, not all macaque visual areas have a clearly defined homologue in the human brain^10^. We therefore examined how effective connectivity is distributed across individual areas by quantifying the percentage of significant responses across all tested connections between early visual areas and the three streams and across the streams themselves. We summarized the results using connectograms (Fig 6) and a stimulation-derived effective connectivity matrix (Fig 7). These representations revealed substantial heterogeneity and directional asymmetry at the area-by-area level (Fig 7): for example, V2 stimulation produced significant responses in 16 of 18 tested connections to IPS0 (88.9%), whereas IPS0 stimulation produced no significant responses in 11 tested connections to V2 (0%).

**Fig 6.**
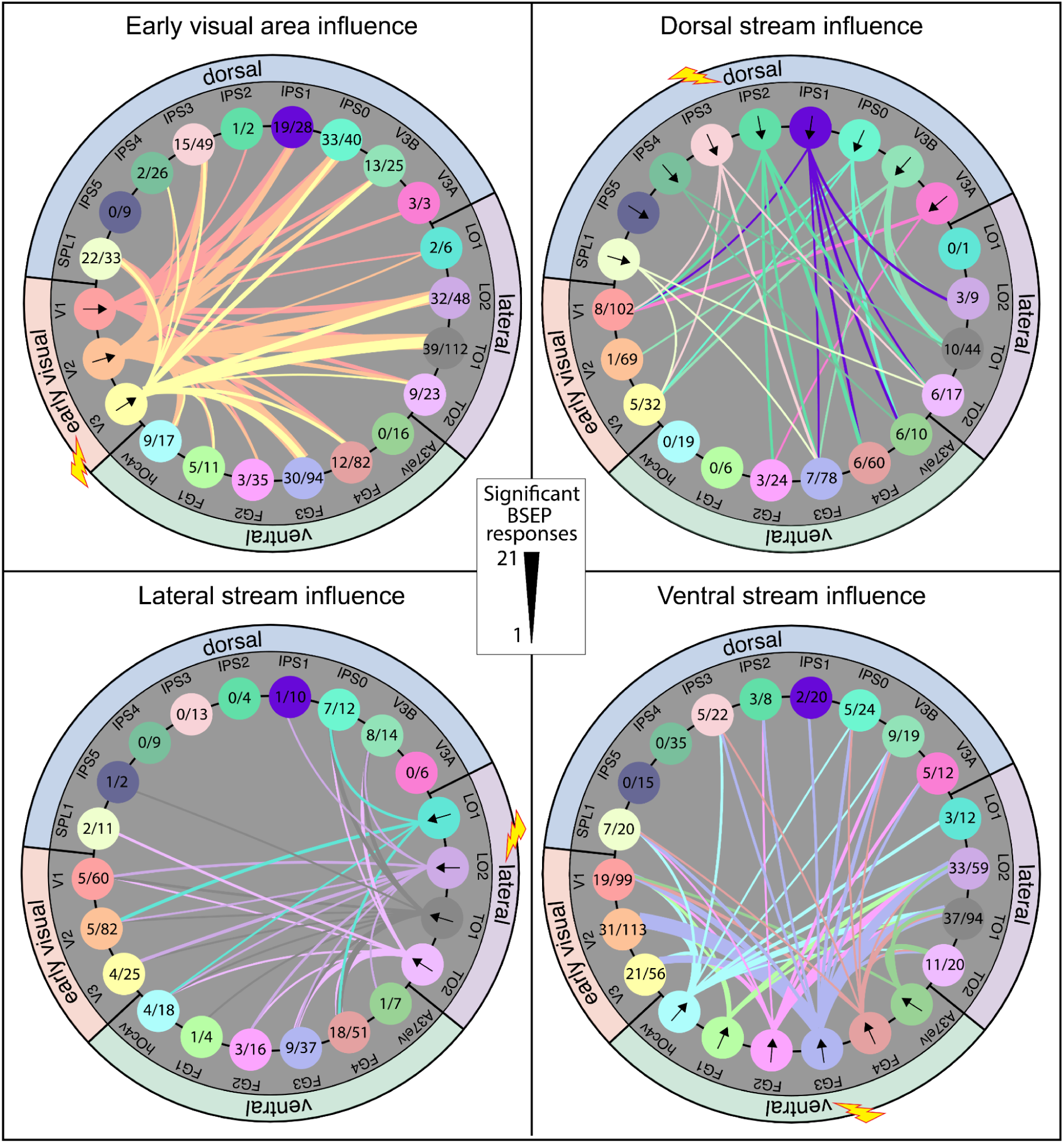
Area-specific organization of effective connectivity across the human visual cortex. Connectograms summarize outgoing effective connectivity from each stimulated visual stream or early visual area group. Each panel shows significant stimulation-evoked influences originating from the group indicated by the lightning bolt icon and terminating in recorded visual areas outside that group. Nodes represent individual visual areas arranged into early visual areas, and the dorsal, lateral, and ventral streams. For recorded areas, the fraction within each node indicates the number of significant incoming responses divided by the total number of tested responses from the stimulated group; arrows within nodes identify areas belonging to the stimulated group. Connecting lines represent significant outgoing influences, with line color indicating the origin area, black arrows denoting direction, and line thickness proportional to the number of significant responses across participants. When stimulation electrode pairs include two adjacent cortical areas, the stimulated pair was counted in both regions, as electrical stimulation affects tissue surrounding both electrode contacts and attribution to a single anode or cathode is not possible^41^. Only tested interactions between the stimulated group and the other visual groups are summarized; within-group interactions are not shown.

**Fig 7.**
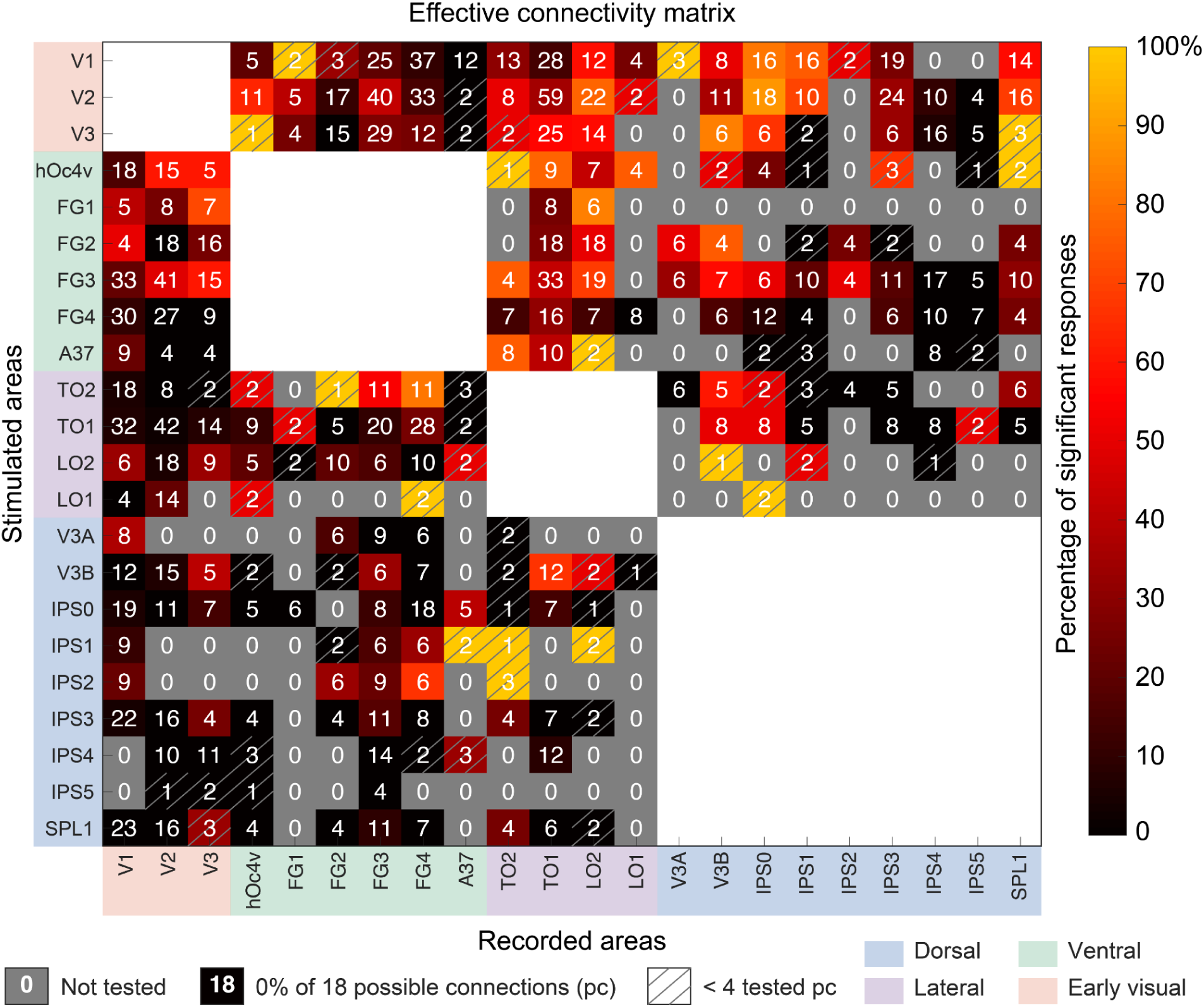
Effective connectivity matrix summarizing stimulation-evoked interactions across visual cortical areas. The matrix shows the percentage of significant stimulation-evoked responses for each stimulation-recording test, calculated across all possible connections given the available electrode coverage. Rows correspond to stimulated areas and columns to recorded areas. Cell colors indicate the percentage of significant responses, and numbers in white indicate the number of tested possible connections (pc). Gray cells denote no tested connection. Hatched cells indicate comparisons with fewer than four tested possible connections and should therefore be interpreted cautiously. Within-group interactions are not shown. This matrix provides a descriptive summary of the empirical pattern of effective connectivity revealed by SPES, highlighting heterogeneous and asymmetric interaction profiles across visual cortical areas. The data and code used to generate this matrix are available as described in the Data and Code Availability sections.

### Dorsal influences converge on A37elv, a ventral temporal region with word-related responses

Connectivity patterns have been linked to functional specialization within the ventral temporal cortex^34^. This relationship supports the hypothesis that interareal connections help shape local functional specialization. Although the overall cross-stream analysis favored temporal-to-parietal influence, the area-level matrix also revealed localized influences in the reverse direction. A particularly consistent example involved stimulation of dorsal stream sites within the intraparietal sulcus (IPS), which elicited significant responses in electrodes on or bordering A37elv, a focal ventral region located in the extreme lateroventral portion of Brodmann area 37 in the posterior occipitotemporal sulcus according to the Brainnetome atlas (Fig 8a). All participants with IPS → A37elv coverage (4 of 4) showed at least one significant evoked response in A37elv. Across these participants, 6 of 10 tested dorsal-to-A37elv connections showed significant responses. Left-hemisphere responsive contacts overlapped or bordered the probabilistic pOTS-characters region^45^, and homologous responses were also observed in the right hemisphere. This overlap motivated us to test whether A37elv showed word-related functional responses.

**Fig 8.**
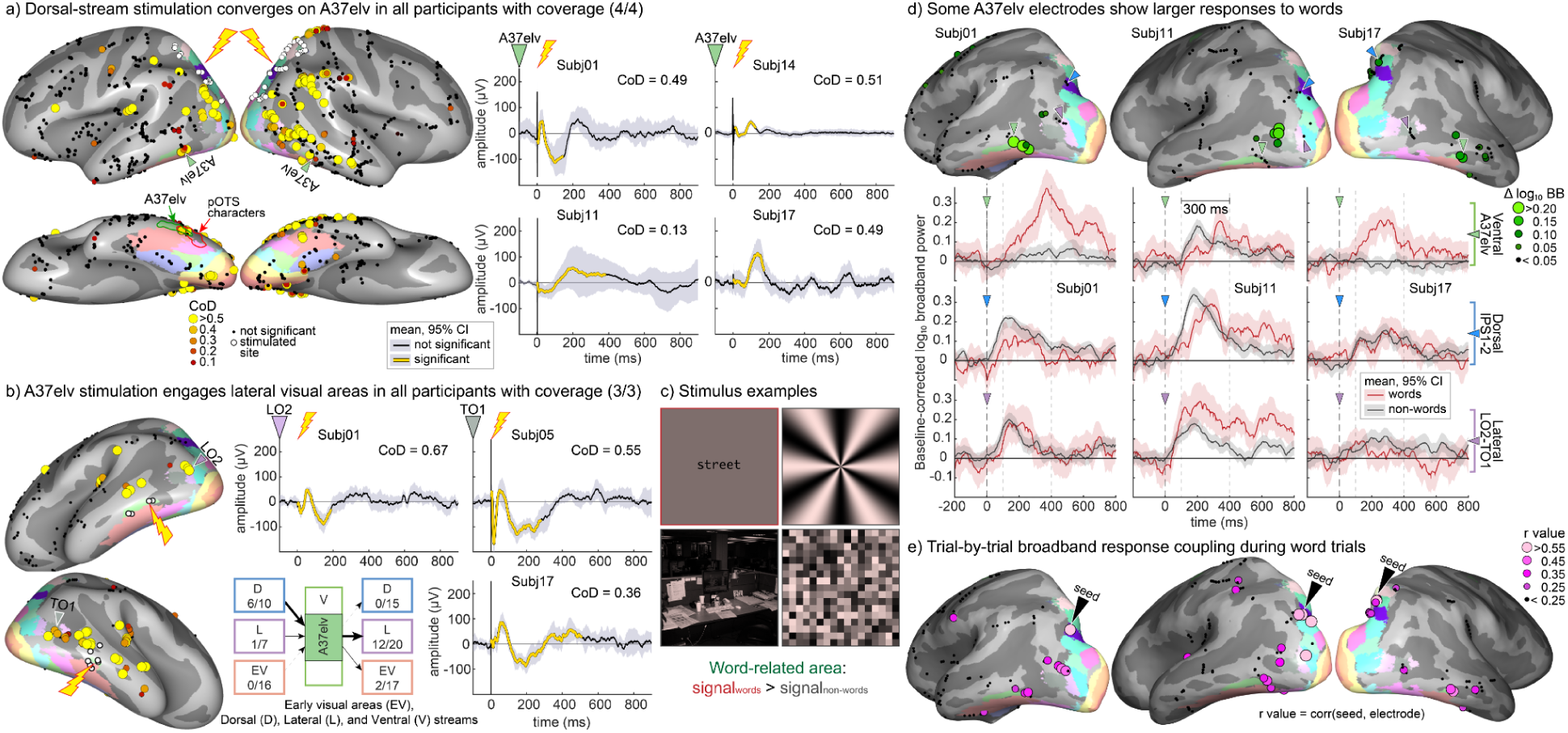
Brodmann area 37 (A37elv) shows selective causal interactions and exploratory word-related responses. **a)** Dorsal stream stimulation evoked significant responses that converged on a focal ventral region located in the extreme lateroventral portion of Brodmann area 37 (A37elv) in the posterior occipitotemporal sulcus. Inflated cortical surfaces show the spatial distribution of significant responses across four participants with dorsal-to-A37elv coverage. Colored dots indicate significant responses, with color denoting coefficient of determination (CoD) values; black dots indicate non-significant recording sites; white circles mark stimulated contact pairs; and the red outline indicates the probabilistic pOTS-characters region^45^ located in the posterior occipitotemporal sulcus. Representative average brain stimulation-evoked responses (BSEPs) recorded in A37elv are shown for each participant ± 95% confidence interval (CI). **b)** A37elv stimulation elicited significant responses primarily in lateral visual areas. Inflated cortical surfaces and representative BSEPs are shown for all three participants with A37elv stimulation and lateral stream coverage. The summary diagram indicates the number of significant responses relative to all tested possible connections for each target stream. All three participants with A37elv-to-lateral coverage showed at least one significant response in lateral visual areas, with 12 of 20 tested connections classified as significant. No significant responses were detected in the sampled dorsal stream (0/15), whereas 2 of 17 tested connections to early visual areas were significant. **c)** Examples of word and other image stimuli used in the visual task. Other images included scenes, gratings, and noise patterns. **d)** Word-related broadband responses in A37elv and other sampled visual areas.

Baseline-normalized log₁₀ broadband responses are shown for word images (red) and other images (gray) in A37elv, dorsal IPS, and lateral LO/TO electrodes from three participants. Broadband power was log₁₀-transformed and baseline-corrected within each run by subtracting the mean log₁₀ broadband power from −200 to 0 ms across all trials in that run (*BB*). Condition-specific traces were calculated as 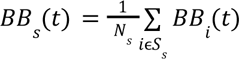, *s* ɛ {*words*, *ot*ℎ*er images*}, where *S_i_* denotes the set of trials in condition *s*, and *N_i_* is the number of trials in that condition. Lines and shaded regions indicate the mean and 95% CI across trials. Colored triangles identify the electrodes whose time courses are shown. Green circles on the inflated cortical surfaces indicate the word-related broadband response difference, with larger circles indicating a larger difference, calculated from the unsmoothed normalized responses as Δ*BB* = *mean*_100−400*ms*_(*BBwords* − *BBot*ℎ*er*). **e)** Exploratory trial-by-trial broadband-response similarity with an IPS seed during word trials. For each word trial, baseline-normalized log₁₀ broadband power was averaged from 10 to 800 ms after image onset for the IPS seed and each electrode. This window summarized trial-level response magnitude rather than temporally precise coupling. Pearson correlations were calculated across word trials as 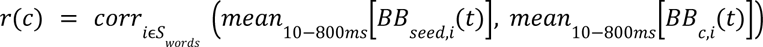 Circle color and size indicate the resulting correlation coefficient. This analysis is descriptive and non-directional and does not establish word-specific coupling or causal influence. Rather, it shows correlations in trial-to-trial response variability during word trials.

Sampling of outputs from A37elv was available in three participants (subjects 01, 05, and 17; Fig 8b). Direct bidirectional comparison of the same IPS and A37elv contacts was not available across all participants. A37elv stimulation elicited significant responses in the lateral stream (LO and TO; Figs 7 and 8b), with at least one lateral response in all participants (3 of 3) with A37elv-to-lateral coverage, and 12 of 20 tested connections were significant. In contrast, A37elv stimulation produced no significant responses in the sampled dorsal stream (0/15), whereas 2 of 17 tested connections to early visual areas were significant. The observed output pattern was therefore concentrated in lateral visual areas, although it was not exclusive to the lateral stream, and these data do not establish reciprocal A37elv–IPS coupling.

A37elv therefore provided a focal example of dorsal-to-ventral influence within the overall temporal-to-parietal directional bias. Because responsive contacts overlapped or bordered the pOTS-characters region, which has been associated with character and word processing, we investigated task responses in three participants (subjects 01, 11, and 17; Fig 8c-d). Images of scenes, noise patterns, gratings, and words were presented to the participants, and we compared high-frequency broadband responses (70-170 Hz) to word images and other images. In two of three participants, broadband responses in a subset of A37elv electrodes were larger to words than to other images; in the third participant, responses showed little separation. These results suggest that some, but not all, A37elv electrodes may participate in word-related visual processing. Interestingly, dorsal and lateral sites that showed effective connectivity with A37elv responded variably to visual inputs, without a consistent word preference (Fig 8d). Effective connectivity therefore does not directly imply shared visual stimulus selectivity. Rather, effective connectivity between the visual streams may be related to other types of information shared during cognitive tasks^21^. To examine such functional covariation in these 3 patients, we correlated responses during word trials, from 10 to 800 ms after image onset (Fig 8e). The descriptive correlation maps showed trial-to-trial broadband-response similarity between the IPS seed, A37elv, and lateral LO/TO sites. Causal stimulation findings may help constrain potential routes of information flow among areas that also show task-related covariation.

## Discussion

### Characterizing causal interactions between human visual areas

The present study provides a system-level characterization of causal interactions across the human visual cortex. Across 23 participants, we constructed a stimulation-derived effective-connectivity matrix spanning 22 atlas-defined visual areas. Controlled, directed cortical perturbations enabled quantitative measurement of the directionality, response reliability, and spatial distribution of interareal influences. This approach moves beyond correlational descriptions of functional connectivity and provides empirical constraints on how activity can propagate across visual streams. Visual-task data from a subset of the same participants provided functional context for selected SPES-defined interactions.

This matrix is especially valuable in humans because some visual regions lack clear one-to-one homologues in non-human primates^10,46^ and functional interactions cannot be inferred directly from anatomical tracing studies. Just as macaque tracing studies have established principles of anatomical connectivity, the present findings provide a foundation for defining the causal organization of interareal influence in the human visual cortex. Interpretation of findings from single-pulse stimulation during sEEG recordings requires some considerations. Here, causal influence refers to a downstream response elicited by controlled perturbation of a cortical site; it does not imply that the effect is mediated by a direct monosynaptic projection. Controlled perturbation is the central strength of this design: by externally driving a defined cortical site, SPES isolates directionality that cannot be recovered from correlated activity alone. At the same time, stimulation-evoked responses do not distinguish direct monosynaptic from indirect polysynaptic effects^47^, and both local and distributed network interactions may contribute to the observed potentials^35,43,44^. Thus, while some influences may arise from direct cortico-cortical projections, others may be mediated through intermediate visual, association, or subcortical regions.

Estimates of response prevalence depend on preprocessing and the operational definition of a significant response. We therefore used the CRP framework together with a cross-validated classification model trained to apply a fixed, reproducible criterion across participants and visual areas. Adjusted common-average re-referencing was used to capture reproducible, time-locked responses reflecting propagation across cortical networks, thereby emphasizing distributed, synchronous responses across cortical regions^48^. Alternative approaches, such as bipolar referencing combined with broadband spectral measures, may provide more spatially focal estimates of local activation^49^ and may be better suited for characterizing local signal generation and nearby spread within approximately 2 cm of surrounding tissue. Finally, intracranial recordings sample the cortex sparsely, and the resulting area-by-area matrix covers 69% of all possible interareal connections within the defined visual network. Despite these limitations, the results constitute an important step toward a stimulation-derived causal map of interareal influence across the human visual cortex and toward understanding how visual areas influence one another.

### Visual information flow and biologically inspired models of vision

Visual processing is often represented as emerging from computations distributed across cortical streams, with information progressively transformed from lower-order to higher-order visual areas^50^. Classical work on the macaque visual system also emphasized distributed hierarchical organization with multiple processing streams^51^. However, interareal interactions in these graphs often remain schematic because their direction, response reliability, and spatial specificity are difficult to measure. This challenge is especially evident when distant regions share similar representational content despite the absence of direct anatomical pathways^50,52^. The matrix derived here helps constrain such graphs by identifying which sampled cross-stream influences are more reliable, directionally biased, and spatially selective at their source or target sites, giving these graphs a more rigorous empirical basis.

There has been substantial progress in computational models of visual representations, particularly through the use of deep neural networks^53–55^. However, links between computational models and neural data are typically made at the representational level; for example, encoding models establish statistical relationships between activity in artificial units and recorded neural responses. Moreover, the mapping between model architecture and brain architecture is often coarse, for example, between network layers and broad biological visual areas such as hV4/LO versus IT/PhC^56^. Although such correspondences are highly informative, they provide limited constraints on biological connectivity because they do not specify or test particular causal interactions between cortical areas or streams^57,58^. Incorporating the matrix provided here into future models could improve their biological grounding and help bridge computational and experimental approaches. The present findings provide empirical constraints on which sampled interactions are asymmetric, reliable, and spatially selective, thereby moving beyond schematic representations toward biologically grounded connectivity architectures.

### Feedforward-feedback asymmetry is stream dependent

Between early visual areas (V1-V3) and higher-order visual streams, feedforward influences were more reliable and detected across a larger fraction of sampled connections than feedback influences overall. Previous intracranial stimulation studies reported a similar feedforward predominance in a small subset of these areas^39^. EEG-fMRI work also reported stronger propagation from visual areas to posterior parietal and inferior temporal regions than in the reverse direction^59^. Here, we show that this asymmetry was pronounced for the dorsal and lateral streams. In contrast, interactions between early visual areas and the ventral stream were comparatively balanced in both response reliability and sampled response prevalence. The causal organization of the visual system therefore does not follow a uniform feedforward hierarchy but instead depends on the visual stream.

These stream-dependent asymmetries should be interpreted in light of both measurement properties and anatomy. Stimulation-based measures of effective connectivity are shaped by multiple factors, including synaptic gain and conduction efficiency^60^. Anatomical studies in macaque visual cortex have shown that feedback pathways are more numerous, whereas feedforward pathways have greater relative anatomical weight^33,61^. These anatomical properties may differ among visual streams and could contribute to the observed differences in response reliability and prevalence.

Functionally, this organization is consistent with distinct roles for feedforward and feedback pathways. Feedforward projections are thought to support the rapid propagation of stimulus-driven information to higher-order areas^32,62–64^, whereas feedback projections have been implicated in imagery^65^, attention^66^, awareness^67^ and other higher-level modulatory processes^68^. Disruptions in feedback strength have been proposed to contribute to visual disorders such as aphantasia and hallucinations^69,70^. Within this framework, the higher response reliability of feedforward influences in the dorsal and lateral stream may support consistent propagation of stimulus-driven activity, whereas the lower sampled reliability of feedback influences may reflect more spatially or contextually selective modulation. The comparatively balanced interaction between early visual and ventral temporal areas further suggests that feedforward and feedback influences may be more closely matched within this portion of the visual network. How these causal interactions are dynamically weighted during perception, attention, imagery, and visually guided behavior remains an important question.

### Directional asymmetries between visual streams

We also observed directional asymmetries between visual streams. Although intracranial electrode coverage necessarily constrained the sampling of cross-stream connections, the combined analysis revealed an overall temporal-to-parietal bias in both response reliability and sampled response prevalence. This bias was not uniform across stream pairs. Ventral-to-lateral and ventral-to-dorsal influences were significantly more reliable than their corresponding reverse directions, whereas the lateral-to-dorsal comparison was not significant. Response prevalence showed the same broad pattern, favoring ventral-to-lateral and ventral-to-dorsal influence but not lateral-to-dorsal influence. The group-level temporal-to-parietal bias was therefore driven primarily by specific ventral outputs rather than by a uniform directional gradient across all stream interactions.

Similar asymmetries have been reported during task performance. Using intracranial electrophysiological recordings and Granger causality, a recent study found stronger ventral-to-dorsal influence than in the reverse direction during memory-guided actions^24^. These findings are compatible with theoretical frameworks proposing that visual processing involves continuous cross-talk between streams rather than strictly segregated parallel pathways^15,16^. In these models, ventral representations inform dorsal computations supporting visuospatial and action-related processing^17,18^. The present temporal-to-parietal bias suggests preferential routing from ventral temporal regions toward lateral and dorsal regions involved in dynamic visual processing, spatial integration, and visually guided action. Whether this orientation changes with attention, action, imagery, or perceptual task demands remains an important question.

### Anatomical limitations in long-range connectivity may constrain cross-stream influence

Our data show that influences between visual streams were not uniformly distributed across all sampled stimulation and recording sites. Cross-stream influence could be broad or spatially restricted at the source end, the target end, or both. This source-side and target-side spatial selectivity is consistent with anatomical constraints indicating that only a small fraction of pyramidal neurons (11%) project beyond adjacent cortical areas^71^. The human cortical surface spans thousands of square millimeters, whereas the long-range pathways linking distant cortical areas have cross-sectional areas on the order of tens of square millimeters. Under such anatomical constraints, electrical stimulation through two small gray matter electrodes (2 mm diameter each) is expected to directly perturb only a limited cortical volume, thereby engaging selective targets within connected regions.

The source and target selectivity analysis (Fig. 5b) refines this notion of spatial selectivity by separating two forms of spatial spread. A directed influence can be broad at the source end if many stimulation pairs within the source area evoke significant responses, or broad at the target end if many recording electrodes within the target area respond. Conversely, an interaction can be spatially specific because only a small subset of source stimulation pairs is effective, because only a small subset of target electrodes responds, or both. Thus, the same interareal interaction can differ in source-side and target-side specificity. This distinction is important because an aggregate measure of significant responses does not indicate whether spatial selectivity arises from the stimulated source sites, the responsive target sites, or both.

An exception to the temporal to parietal bias was the influence of dorsal IPS stimulation on ventral A37elv. Dorsal stimulation elicited significant A37elv responses in all four participants with relevant coverage. Responsive contacts overlapped or bordered the probabilistic pOTS-character region, a ventral temporal region associated with character and word processing^45,72,73,74^. The ventral temporal cortex is influenced not only by bottom-up visual input but also by task demands and top-down attentional signals, potentially originating from the dorsal visual stream^21,75,76^. Reading depends on rapid eye movements and coordinated visuospatial attention, which require interactions between dorsal attentional systems and ventral visual regions^77,78^. In this context, dorsal influences on A37elv may help coordinate visuospatial demands with ventral visual representations during reading and symbol recognition. The exploratory task data showed word-related broadband-response differences in some sampled A37elv electrodes, but did not determine the functional role of the dorsal-to-A37elv interaction. Direct perturbation during task performance will be required to test this interpretation.

The interaction pattern surrounding A37elv was asymmetric. A37elv stimulation preferentially influenced lateral visual areas while no significant responses were detected in the sampled dorsal stream. Together, these observations are consistent with an asymmetric interaction pattern: dorsal influences reach A37elv, whereas A37elv preferentially influences lateral visual cortex. Similar area-specific asymmetries were observed in the effective-connectivity matrix and are consistent with anatomical observations in non-human primates showing that a substantial proportion of interareal connections are unidirectional^33,79,80^. Task-response profile similarity linked IPS, A37elv, and lateral sites but did not determine direction. Together, these findings suggest that cross-stream communication can be organized through asymmetric and spatially restricted causal interactions.

### Outgoing/incoming asymmetries and organizational profiles within the visual cortical network

We summarized distinct organizational profiles for visual cortical regions by examining the ratio between outgoing and incoming effective connectivity within the sampled visual cortical network, using a similar logic as previous work^42–44^. Early visual areas exhibited a high output-to-input ratio, consistent with a source-like profile in which influence was broadly distributed to the three visual streams. The ventral stream showed a similar but more moderate bias toward outgoing influences, also consistent with a source-like profile within the sampled network, potentially transmitting elaborated visual representations to other regions. In contrast, the lateral and dorsal streams exhibited more incoming than outgoing influence, consistent with integrator-like profiles within the visual cortical network. This organization suggests that cortical visual processing emerges from interactions between source-like and integrator-like regions within a structured network architecture.

## Conclusions

In summary, this study establishes a stimulation-derived effective-connectivity matrix of the human visual cortex, providing a quantitative description of the directionality, response reliability, and spatial distribution of causal interareal influences. Our results reveal a selective and stream-dependent causal architecture with source-like early visual and ventral temporal regions and integrator-like dorsal and lateral regions. Rather than a uniform feedforward hierarchy, the human visual system is organized as a heterogeneous causal network in which the direction and spatial extent of influence depend on the streams and areas involved. These findings define a causal response architecture through which information may flow across the human visual cortex and provide empirical constraints for biologically grounded models of visual computations.

## Methods

### Participants

Twenty-three patients (mean age = 25.9 years, SD = 8.98; 10 males, 13 females) undergoing sEEG for clinical evaluation of epilepsy participated in this study. Stimulation was delivered through selected pairs of neighboring contacts, and BSEPs were recorded from all remaining contacts (Fig 9a). Only participants with gray-matter electrodes located in at least two of the four regions of interest (early visual areas, dorsal stream, lateral stream, or ventral stream) were included (Fig 9b). All participants provided written informed consent prior to participation, and all procedures were approved by the Institutional Review Board of Mayo Clinic (IRB # 15-006530).

**Fig 9.**
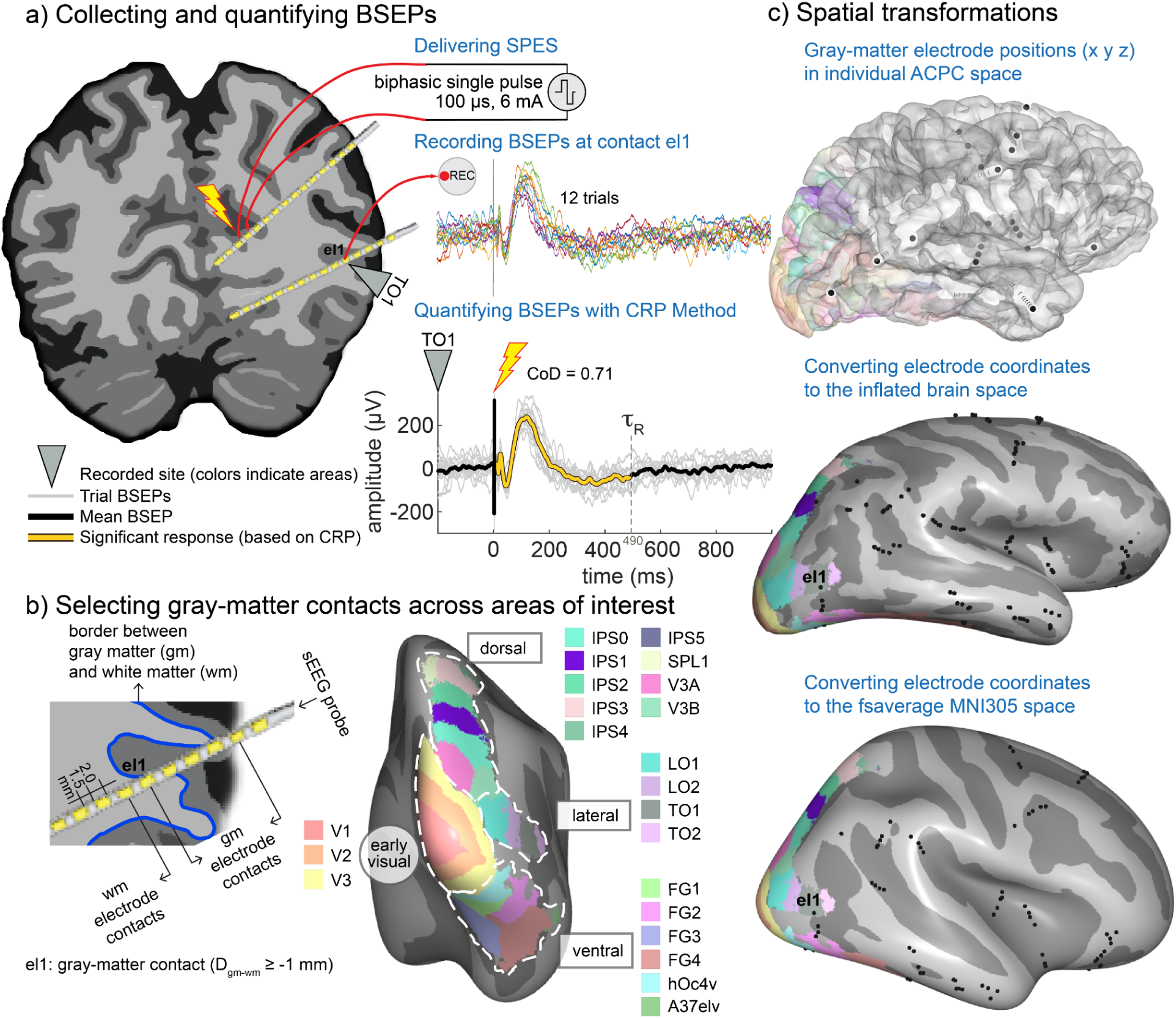
Quantifying effective connectivity between human visual cortical areas. **a)** Electrode pairs in the gray matter were stimulated with ∼12 single electrical pulses (SPES) and brain stimulation-evoked potentials (BSEPs) were recorded simultaneously from all remaining contacts. BSEPs were quantified using the Canonical Response Parameterization (CRP) method, yielding measures of response reliability (coefficient of determination, CoD), response duration (*τ*_R_), signal-to-noise ratio (Vsnr), and a CRP-derived p value (pVal). Responses were classified as significant or non-significant using the fixed classification model described in the Statistical analyses section. The lightning bolt icon marks the stimulation site, and the inverted triangle indicates an example recording contact (el1) located in TO1 of the lateral stream. **b)** Gray-matter contacts across visual areas of interest were selected by calculating the distance between each contact center and the gray-white matter boundary (blue line) in the individual T1-weighted MRI (T1w). Contacts with a distance ≥ -1 mm from the boundary were classified as gray-matter contacts and included in the analyses. Visual cortical areas were defined using probabilistic atlases^13,81–83^ and grouped into four areas of interest (early visual areas; dorsal stream; lateral stream; ventral stream). **c)** Ipsilateral gray-matter electrode coordinates were determined in individual ACPC space from post-implantation CT scans co-registered to preoperative T1w (top), projected onto the individual inflated cortical surface (middle), and transformed to MNI305 space and displayed on the fsaverage surface for group-level visualization (bottom).

### Electrode localization and definition of areas of interest

To obtain electrode positions, post-implantation CT scans were co-registered to preoperative T1-weighted (T1w) MRI volumes using mutual information. Stereotactic coordinates (x, y, z) for each sEEG contact were extracted as previously described^84^. Cortical surface reconstructions were generated using the FreeSurfer recon-all pipeline^85,86^. Individual cortical parcellations were obtained using Neuropythy^81^ (Fig 9b-c). For network-level analyses, we defined a visual cortical network comprising four areas of interest based on probabilistic atlases. Early visual areas consisted of V1, V2, and V3^81^. The dorsal stream comprised IPS0, IPS1, IPS2, IPS3, IPS4, IPS5, SPL1, V3A, and V3B^13^. The lateral stream consisted of LO1, LO2, TO1, and TO2^13^. The ventral stream comprised FG1-FG4, hOc4v^82^, as well as the A37elv region^83^. Each sEEG electrode consisted of multiple cylindrical contacts (typically 2 mm length, with 1.5 mm spacing).

Only ipsilateral electrodes located in gray matter were included in the analyses. This is essential to make inferences about directionality, because white matter stimulation can induce both orthodromic and antidromic responses^41^, see S1 Fig. The distance between each contact center and the FreeSurfer-based estimate of the gray-white matter boundary was calculated, with positive distances indicating gray matter and negative distances indicating white matter (Fig 9b). Contacts with centers at a distance ≥ -1 mm were classified as gray-matter (gm) contacts. A contact was assigned to a visual area if its center lay within 4 mm of any vertex belonging to that area.

### Experimental design and data preprocessing

The experimental design combined causal perturbations via SPES with sEEG recordings to characterize effective connectivity across early visual areas and dorsal, lateral, and ventral visual streams. Directional influences were quantified by sampling all possible connections between the four visual groups (regions of interest) within the cortical visual network, where a possible connection was defined as a combination of a stimulated electrode pair and a recording electrode for which effective connectivity could be measured.

### Electrical stimulation

During stimulation, patients were awake and resting with open eyes (e.g., viewing a screen or conversing). SPES was delivered through pairs of neighboring contacts using biphasic square-wave pulses (6 mA amplitude, 100 or 200 µs width per phase, ∼12 trials per stimulation site, and a 2-5 s interstimulus interval). These parameters are consistent with prior work^87,88^ demonstrating reliable cortical responses and network-level propagation following single-pulse stimulation in humans. In particular, higher current intensities (> 5 mA) combined with short pulse widths per phase (<0.3 ms) are associated with more robust stimulation-evoked response metrics^88^. These stimulation parameters have been shown to reliably elicit local and distant cortical responses and network-level propagation in human intracranial studies^35,87–89^, supporting their use for probing cortico-cortical responses and effective connectivity.

Two stimulation-recording systems were used: a g.Estim stimulator with a g.HIamp amplifier (100 µs per phase; 4800 Hz sampling rate; g.tec, Schiedlberg, Austria) and a Nicolet cortical stimulator with a Natus Quantum amplifier (100-200 µs per phase; 2048 Hz sampling rate; Natus, Wayzata, MN, USA). BSEPs were recorded simultaneously from all remaining contacts.

### Exploratory visual task and broadband response analyses

Three participants (subj01, subj11, and subj17) completed a visual task using synthetic images that contained natural scenes, words, noise patterns, and gratings from the synthetic Natural Scenes Dataset (NSD-synthetic^90^, Fig. 8c). Broadband power (70–170 Hz) was log₁₀-transformed and baseline-corrected separately within each run by subtracting the mean log₁₀ broadband power from −200 to 0 ms across all trials in that run. The resulting normalized log broadband power was analyzed separately for word images and other images, and time courses were displayed as means with 95% confidence intervals. Word-related response magnitude was calculated for each electrode as ΔBB = BB_words_ − BB_other_. Because task data were available for only three participants, these analyses were considered descriptive and exploratory, and no across-participant inferential claim was made.

To examine functional similarity among connected sites, baseline-normalized broadband responses were averaged from 10 to 800 ms after image onset for each word trial. We selected this extended interval to capture word processing-related variability. Within each participant, the resulting trial-response vector from a dorsal IPS seed electrode, selected because stimulation at that site produced an A37elv BSEP during SPES, was correlated with the corresponding vector at every other bipolar electrode using Pearson’s correlation coefficient across the 40 word trials. The maps were descriptive. This analysis measured non-directional similarity in trial-to-trial response magnitude during word trials.

### Data preprocessing

Channels that contained artifacts upon visual inspection or that were located outside the brain or within seizure onset zones were excluded and marked as bad. For each stimulation pair, recording contacts located within 5.5 mm of either stimulated contact were excluded from that possible connection. Adjusted common-average re-referencing, using only the 25% of channels with the lowest variance to avoid introducing bias, was applied within each 64-channel amplifier, excluding channels previously marked as bad^91^. Continuous recordings were segmented from -2.5 to 2.5 s relative to stimulation onset (0 ms), and baseline correction was applied using the -100 to -10 ms window. Residual stimulation-related artifacts were identified by fitting an exponential-plus-offset function to the baseline-corrected evoked response within a 2.5–80 ms post-stimulation window. Responses with R² ≥ 0.80 and a fitted decay time constant of 1–20 ms were flagged as artifact candidates. Only artifact-free responses from gray-matter electrodes in the same hemisphere as the stimulation site, and within predefined visual regions were included in the analyses. The spatial distribution of evoked responses was inconsistent with volume conduction as the primary explanation for the observed effects. Volume conduction would be expected to produce the largest responses near the stimulation site, followed by a gradual decay with distance^92,93^. The measured responses did not show this expected distance-dependent decay and instead exhibited spatial selectivity more consistent with network-mediated propagation. Instead, evoked response features were consistent with network-level propagation of stimulation-evoked activity. For visualization, responses were displayed on inflated cortical surfaces^94^.

### Statistical analyses

#### Evoked response significance and reliability

Brain stimulation-evoked potentials were characterized using Canonical Response Parameterization (CRP^37^), which parameterizes stimulation-evoked responses and provides waveform-independent measures of response duration, strength, and reproducibility. Analyses were performed starting after the stimulation artifact (3 ms in the g.tec system and 11 ms in the Natus system) to 1 s post-stimulation. For each possible connection, CRP-derived features included signal-to-noise ratio (Vsnr), p-value (pVal), the duration of the response (*τ*_R_), explained variance, summaries of the trial projection coefficients (α′), and CRP t and p statistics. The coefficient of determination (CoD) was used to quantify cross-trial response reliability.

As a measure of cross-trial response reliability, we calculated the coefficient of determination (CoD):

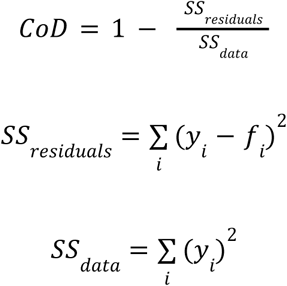

where *y_i_* is the measured response amplitude and *f_i_* is the predicted response amplitude for trial *i*.

The total number of possible connections in each direction varied across participants depending on electrode number and placement. To classify responses consistently across participants and visual areas, we trained a logistic-regression model on expert-reviewed response labels from 11 participants using CRP-derived features and CoD. Model performance was evaluated using leave-one-participant-out cross-validation. The selected model and probability threshold (0.30) were then fixed and applied to the complete cohort. Responses with probabilities ≥ 0.30 were classified as significant, and artifact rejection was performed separately.

#### Response reliability and sampled response prevalence

*Response reliability* was quantified using CoD across possible connections in a given direction. Higher CoD values indicate more reproducible stimulation-evoked responses across trials. *Sampled response prevalence* was defined as the proportion of responses classified as significant among all possible connections. A high proportion indicates detectable influence across a large fraction of stimulation-recording combinations, whereas a low proportion indicates detectable influence across a smaller fraction of these combinations. Source and target ratios were used to determine whether spatial extent or selectivity arose at the stimulated or responsive sites.

#### Output-to-input ratio

To characterize the relative role of each area of interest within the defined visual cortical network, we quantified the ratio between outgoing and incoming effective connectivity. For each region, we computed an output-to-input ratio defined as the proportion of significant outgoing connections divided by the proportion of significant incoming connections. While interpretation requires caution because of sampling restrictions, this ratio provides an estimate of whether a region exhibits a source-like profile (ratio > 1) or an integrator-like profile (ratio < 1) within the sampled network^42–44^.

#### Source and target spatial influence

To quantify the spatial spread of each directed interregional influence, we separately estimated source and target ratios for each directed source-target interaction. Source stimulation pairs were defined as unique participant-by-stimulation-pair combinations, and target electrodes were defined as unique participant-by-recording-electrode combinations. The source ratio was defined as the percentage of source stimulation pairs that evoked at least one significant response in the target region. The target ratio was defined as the percentage of target recording electrodes that showed at least one significant response to stimulation from the source region. Thus, the source ratio captures how broadly effective stimulation sites were distributed within the source region, whereas the target ratio captures how broadly significant responses were distributed within the target region.

#### Linear mixed effects models

Directional differences in response reliability were tested using linear mixed-effects models implemented in MATLAB (fitlme) and fitted using maximum likelihood (ML):

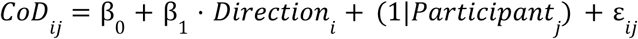

where *CoDij* is the coefficient of determination for possible connection *i* from participant *j*, β0 is the intercept, β1 captures the fixed effect of Direction, (1|*Participanti*) denotes a participant-specific random intercept, and ε*ij* is the residual error term. Direction was coded so that positive β1 values favor feedforward or temporal-to-parietal influence.

Two primary planned comparisons were performed using independent models: (1) feedforward direction (early visual areas → visual streams) versus feedback direction (visual streams → early visual areas); and (2) temporal-to-parietal direction (ventral → lateral, ventral → dorsal, and lateral → dorsal) versus parietal-to-temporal direction (dorsal → lateral, dorsal → ventral, lateral → ventral). Additional pairwise comparisons were performed within each analysis to test whether these effects were consistent across specific streams. For the feedforward-feedback analysis, comparisons were conducted separately for dorsal, lateral, and ventral streams. For the temporal-to-parietal analysis, comparisons were conducted separately for ventral-lateral, ventral-dorsal, and lateral-dorsal streams. The unit of observation was the possible connection.

Because only two primary a priori hypotheses were tested, no additional correction for multiple comparisons was applied to the primary analyses. These primary comparisons are reported using unadjusted p values. Within each analysis, false discovery rate (FDR) correction was applied jointly to the three follow-up pairwise comparisons, which are reported as q values. Model parameters and associated statistics (β, SE, t, and p; and q for the FDR-corrected comparisons) are reported in S1 and S2 Tables.

#### Visualizing effective connectivity

To visualize group-level results, the electrode coordinates were previously transformed to MNI305 space and displayed on the fsaverage surface (Fig 9c), creating a compound map that includes the corresponding CoD values. This visualization summarizes the spatial distribution and relative reliability of outgoing effective connectivity following stimulation of individual electrode pairs. Circular connectograms and the effective-connectivity matrix were generated in MATLAB to illustrate the organization of effective connectivity.

## Data availability

The complete dataset required to reproduce all results will be made publicly available upon manuscript acceptance.

## Code availability

All analyses were implemented in MATLAB R2024a. Analysis code, including the code used to generate the effective-connectivity matrix is publicly available at https://github.com/MultimodalNeuroimagingLab/visual_connectivity.

## Acknowledgements

We thank the patients for participation in this study; and Cindy Nelson, Karla Crockett, and other staff at Saint Mary Hospital, Mayo Clinic, Rochester, MN for assistance and Dr. Hiromasa Takemura for discussions.

## Supporting information

**S1 Fig.**
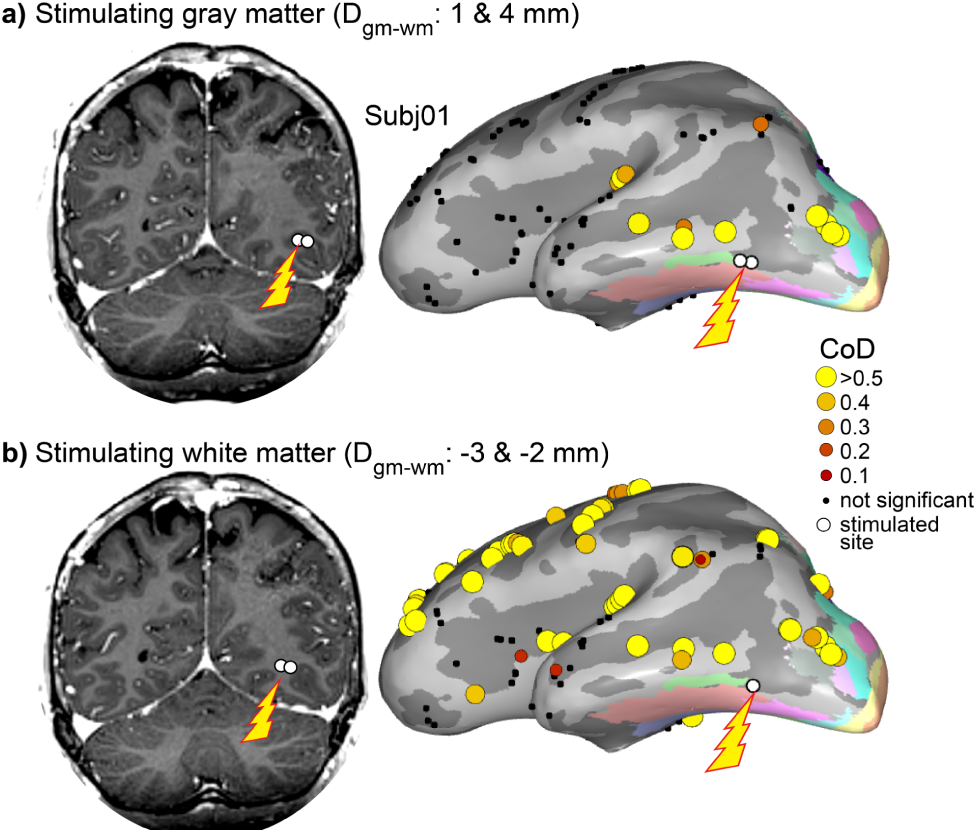
White-matter stimulation produces widespread responses compared to gray-matter stimulation. Representative example from one participant (subj01). **a)** Stimulation of an electrode pair located in gray matter (distance from the gray matter boundary, D_gm-wm_: 1 and 4 mm) elicited spatially localized responses with higher CoD values at a limited set of recording sites. Left: coronal T1-weighted slice showing the stimulated electrode pair (white circles; lightning bolt). Right: responses projected onto the inflated cortical surface. **b)** In contrast, stimulation of an electrode pair located in white matter (D_gm-wm_: -3 and -2 mm) elicited more widespread responses across distributed sites. This white matter stimulation may directly activate portions of major white-matter pathways, such as the arcuate fasciculus and the vertical occipital fasciculus, which could contribute to the responses observed in frontal and parietal regions. These examples illustrate that white-matter stimulation is more likely to induce broad and less spatially specific responses, consistent with current spread along fiber tracts and mixed orthodromic and antidromic propagation. To ensure spatially specific and interpretable estimates of effective connectivity directionality, only gray-matter electrode contacts were used for stimulation and recording in the main analyses.

**S1 Table.** Feedforward vs Feedback response reliability models. Linear mixed-effects model results for comparisons of CoD response reliability between feedforward and feedback directions, including follow-up comparisons for early visual areas (EV), dorsal (Do), lateral (La), and ventral (Ve) streams. The ‘group’ term denotes the fixed effect of direction. Estimates are reported as β_1_ ± SE. DF = degrees of freedom; CI = confidence interval. Follow-up pairwise comparisons for the dorsal and lateral streams remained significant after false discovery rate correction, whereas the ventral comparison was not significant.

| Comparison | Effect | $\beta \pm SE$ | t | DF | p | q | 95% CI |
| --- | --- | --- | --- | --- | --- | --- | --- |
| Feedforward vs Feedback | Intercept | 0.12322 ± 0.024824 | 4.9638 | 1094 | 8.01E-07 | | [0.074514, 0.17193] |
| | group | 0.13237 ± 0.01399 | 9.4615 | 1094 | 1.79E-20 | | [0.10492, 0.15982] |
| EV→Do vs Do→EV | Intercept | 0.066039 ± 0.050934 | 1.2966 | 337 | 0.19567 | | [-0.034149, 0.16623] |
| | group | 0.21252 ± 0.023788 | 8.9342 | 337 | 2.73E-17 | 8.19E-17 | [0.16573, 0.25931] |
| EV→La vs La→EV | Intercept | 0.10295 ± 0.033185 | 3.1023 | 309 | 0.0020977 | | [0.037652, 0.16825] |
| | group | 0.19066 ± 0.024088 | 7.9151 | 309 | 4.46E-14 | 6.69E-14 | [0.14326, 0.23806] |
| EV→Ve vs Ve→EV | Intercept | 0.17067 ± 0.040709 | 4.1923 | 444 | 3.33E-05 | | [0.09066, 0.25067] |
| | group | 0.038541 ± 0.021445 | 1.7972 | 444 | 0.072977 | 0.073 | [-0.0036045, 0.080686] |

**S2 Table.**
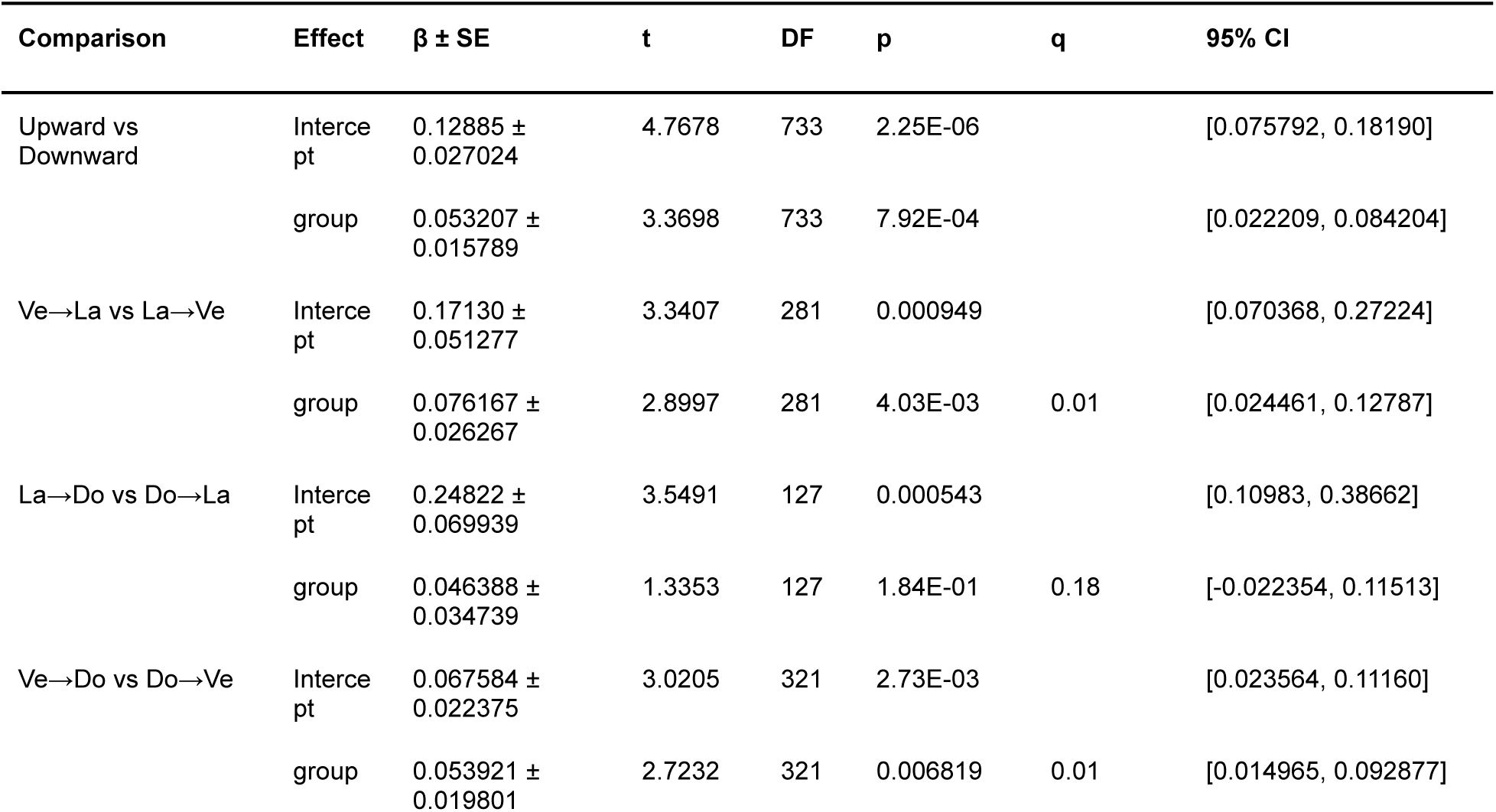
Upward vs Downward response reliability models. Linear mixed-effects model results for comparisons of CoD response reliability between temporal-to-parietal (upward) and parietal-to-temporal (downward) directions, including follow-up comparisons for dorsal (Do), ventral (Ve), and lateral (La) streams. The ‘group’ term denotes the fixed effect of direction in each comparison. Estimates are reported as β_1_ ± SE. DF = degrees of freedom; CI = confidence interval. The combined comparison across the three stream pairs was significant; after false discovery rate correction, the ventral–lateral and ventral–dorsal comparisons were significant, whereas the lateral–dorsal comparison was not.

